# clusterProfiler: An universal enrichment tool for functional and comparative study

**DOI:** 10.1101/256784

**Authors:** Guangchuang Yu

## Abstract

I present the open-source software package clusterProfiler (https://github.com/GuangchuangYu/clusterProfiler) for functional enrichment analysis. clusterProfiler provides a universal interface for functional enrichment analysis for internal supported ontologies/pathways as well as annotation data provided by users or obtained from online databases. It enables comparative analysis and offers comprehensive visualization tools for result interpretation.

---

Functional enrichment analysis is one of the most widely used technique for interpreting gene lists or genome-wide regions of interest (ROIs) derived from high-throughput sequencing (HTS). Although tools are proliferating to perform gene-centric or epigenomic enrichment analysis, most of them are designed for model organisms or specific domains (*e.g.* fungi^1^ or plant^2^ *etc*.) embedded with particular annotations (*e.g.* Gene Ontology (GO) or Kyoto Encyclopaedia of Genes and Genomes (KEGG)). Non-model organisms and functional annotations other than GO and KEGG are poorly supported. Moreover, an increasing concern upon the quality of gene annotation has raised an alarm in biomedical research. A previous study^3^ reported that about 42% of the tools were outdated by more than five years and functional significance were severely underestimated with only 26% of biological processes or pathways were captured compare to using up-to-date annotation. Reanalysing GTEx dataset^4^ published by ENCODE consortium using clusterProfiler uncovered a lot of new pathways that leads to generate new hypotheses (https://github.com/GuangchuangYu/enrichment4GTEx_clusterProfiler). Such negative impacts of outdated annotation can be propagated for years and hinder follow-up studies.

The clusterProfiler package was designed by considering the supports of multiple ontology/pathway annotations, up-to-date gene annotations, multiple organisms, user’s annotation data and comparative analysis. The package employs modular design and supports disease analyses (Disease Ontology^5^, Network of Cancer Gene^6^, gene-disease and variant-disease associations^7^), Reactome pathway and Medical Subject Headings (MeSH) analysis via its sub-packages DOSE^5^, ReactomePA^8^ and meshes (https://github.com/GuangchuangYu/meshes). The clusterProfiler package internally supports GO and KEGG. GO annotation data can be obtained directly from Bioconductor OrgDb packages, or retrieved from web resources (*e.g.* AnnotationHub, biomaRt^9^ and UniProt^10^ *etc*.). KEGG Pathway and Module were directly obtained using KEGG API and supports more than five thousand genomes (http://www.genome.jp/kegg/catalog/org_list.html) as well as KEGG Orthology (KO) which is particular useful for metagenomic functional studies. In addition, clusterProfiler provides general functions, enricher and GSEA, to perform fisher’s exact test and gene set enrichment analysis using user defined annotations, making it possible to support novel functional annotation of newly sequenced species (*e.g.* electronic annotation using Blast2GO^11^ and KAAS^12^), unsupported ontologies/pathways (*e.g.* InterPro Domain, Clusters of Orthologous Groups, Mouse phenotype ontology *etc*.) or customized annotation. clusterProfiler provides parser functions to import GMT file so that gene sets downloaded from Molecular Signatures Database can be directly supported as well as to retrieve whole genome GO annotations from UniProt database using taxonomic ID. Epigenomic enrichment analysis is also supported with user-provided ROIs in BED format by combining with in-house annotation package ChIPseeker^13^. GO semantic similarity measurement implemented in another in-house package GOSemSim^14^, which allows calculating GO term similarity using several methods based on information content and graph structure, was also incorporated in clusterProfiler to remove redundancy of enriched GO terms. This feature simplifies result and assists in interpretation as well as against annotation/interpretation bias.

Albeit functional enrichment analysis is commonly used in downstream analysis to decipher key biological processes, visualization of enriched result is mainly depicted by bar chart of most significant terms, which maybe dominated by redundant terms and cannot reveal the complex of gene-pathway associations. The clusterProfiler sub-package, enrichplot (https://github.com/GuangchuangYu/enrichplot), is designed to visualize enrichment result by integrating expression data (Fig. 1). These methods allow users without programming skills to generate effective visualization to explore and interpret results as well as to find patterns within the data. In addition, all these visualization methods were implemented based on ggplot2, which allows customization using grammar of graphics.

**Figure 1.**
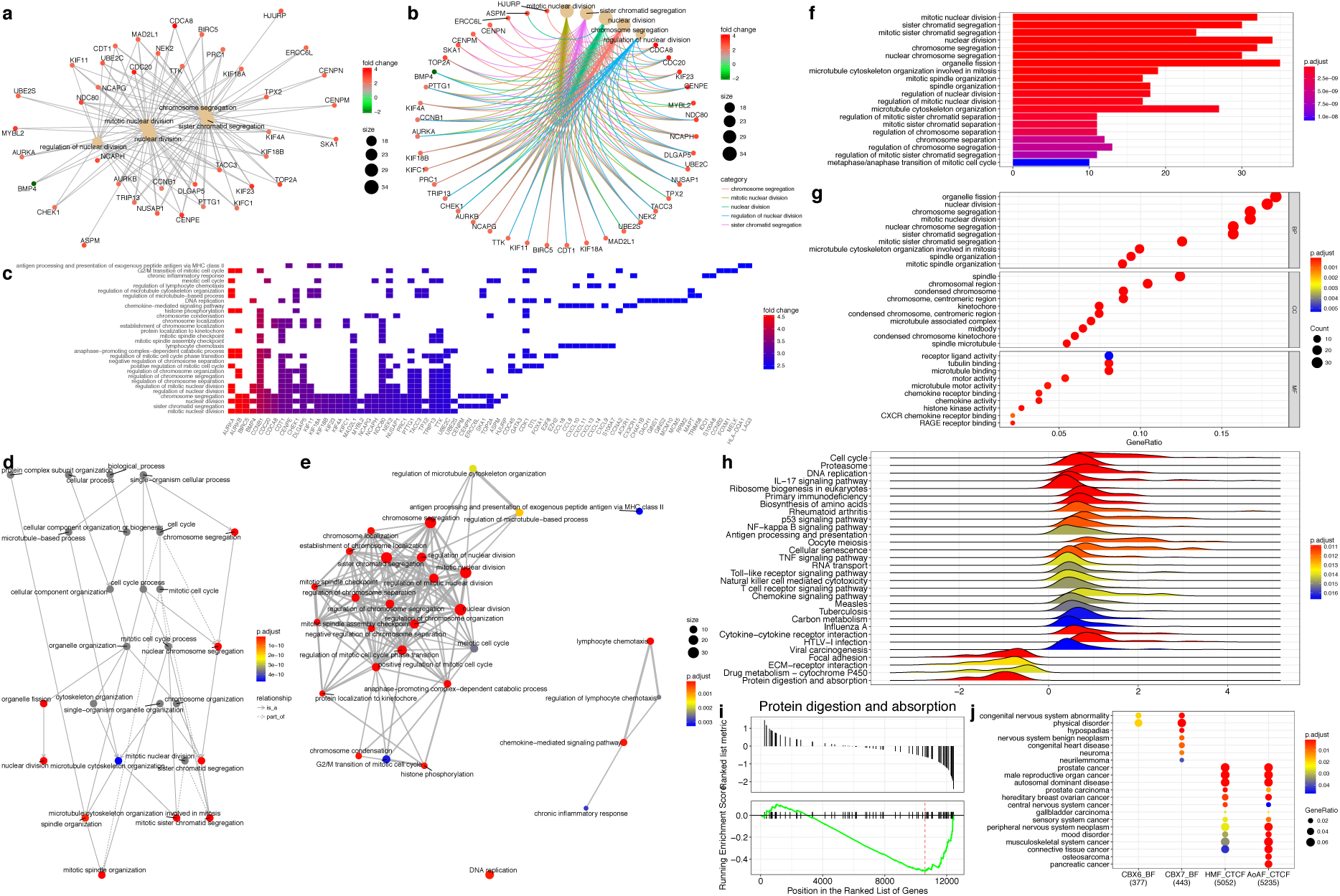
Visualization methods for enrichment result. (a) Gene-concept network depicts the linkages of genes and biological concepts as a network. (b) Circular version of gene-concept network. (c) Displaying gene-concept relationship as a heatmap with gene expression data integrated. (d) Directed acyclic graph induced by most significant GO terms. (e) Enrichment map organizes enriched terms into a network with edges weighted by the ratio of overlapping gene sets. Mutually overlapping gene sets are tending to cluster together, making it easy to identify functional modules. (f) Bar chart to display gene count or ratio as bar height and coloured by enrichment scores (e.g. p.adjust). (g) Dot chart that similar to bar chart with capability to encode another score as dot size (also known as bubble plot). Both bar and dot charts support faceting to visualize sub-ontologies simultaneously. (h) Ridge line plot for expression distribution of GSEA result. By default, only display the distribution of core enriched genes (also known as leading edge) and can be switched to visualize the distribution of whole gene sets. (i) Running score and pre-ranked list of GSEA result. (j) Comparing enriched results across multiple experiments. This example compared disease associations of multiple genome-wide ROIs (data from GSM1295076, GSM1295077, GSM749665, GSM749666).

Comparing functional profiles reflects functional consensus and difference among different experiments and helps identifying differential functional modules in omic datasets. With the infrastructure of clusterProfiler to support a wide range of ontology/pathway annotations and plenty of organisms, the comparison can be applied to many circumstances based on gene or ROI levels. Furthermore, the compareCluster function supports using formula, which is widely used in R language to express statistical model, making it easy to compare functional profiles for different conditions and at different time points.

## AUTHOR CONTRIBUTIONS

G. Y. conceived and designed this project, implemented the algorithms and software packages and wrote the manuscript.

## COMPLETING FINANCIAL INTERESTS

The author declares no competing financial interests.

